# *In vivo* reprogramming of wound-resident cells generates skin with hair

**DOI:** 10.1101/2023.03.05.531138

**Authors:** Yuta Moriwaki, Shen Qi, Hiroyuki Okada, Du Zening, Shogo Suga, Motoi Kato, Takao Numahata, Kexin Li, Koji Kanayama, Mutsumi Okazaki, Yusuke Hirabayashi, Juan Carlos Izpisua Belmonte, Hironori Hojo, Masakazu Kurita

## Abstract

Mammalian skin appendages, such as hair follicles and sweat glands, are complex mini-organs formed during skin development^1, 2^. As wounds heal, the resulting scar tissue lacks skin appendages. The clinical regeneration of skin appendages is an ongoing challenge^3, 4^. Skin epithelial tissues have been regenerated *in vivo* by cellular reprogramming^5, 6^, but the *de novo* generation of skin appendages has not previously been achieved. Here, we show that transplantation of a type of epithelial cell and two types of mesenchymal cells, reprogrammed from adult mouse subcutaneous mesenchymal cells to mimic developing skin cells, resulted in the generation of skin-appendage-like structures. Furthermore, with the development of a new AAV serotype, *in vivo* reprogramming of wound-resident cells with the same reprogramming factors generates skin with *de novo* appendages in adult mice. These findings may provide new therapeutic avenues for skin regeneration and frequent aging-associated skin appendage disorders, such as hair loss and dry skin, and may extend to other tissues and organs. This study also provides the potential for *de novo* generation of complex organs *in vivo*.

## Main

After recent advances in cellular reprogramming^5, 6^, we have developed a method to generate skin epithelial tissues by *in vivo* reprogramming of wound-resident mesenchymal cells with four transcription factors (*DNP63A, GRHL2, TFAP2A,* and *cMYC*), resulting in cells with the ability to form stratified epithelia, which we call induced stratified epithelium progenitors (DGTM-iSEPs)^7^. *De novo* epithelialization can be induced from the surface of an ulcer^8^, but no skin appendages were present in the regenerating skin. As skin appendages form during skin development^1, 2^, we hypothesized that wound-resident adult mesenchymal cells could be reprogrammed to epithelial and mesenchymal cells similar to that of developing skin, thus generating skin with appendages *in situ*.

### Skin reconstitution assay

To identify sets of direct reprogramming genes to generate cells capable of regenerating skin appendages, a traditional skin reconstitution assay^9, 10^ was used as the functional test. While the transplantation of mixtures of adult skin-derived epithelial cells (ASECs) and adult subcutaneous mesenchymal cells (ASMCs) into a silicone chamber attached to a skin ulcer generated on the back of immunodeficient mice^11^ resulted in no skin appendage regeneration (Fig. 1a), skin cells from E14.5 foetuses (embryonic skin cells, ESKCs) robustly regenerated skin appendages (Fig. 1b). Mixtures of neonatal skin epithelial cells (NSECs) and neonatal skin mesenchymal cells (NSMCs) resulted in moderately regenerated skin appendages (Fig. 1c).

**Fig. 1:**
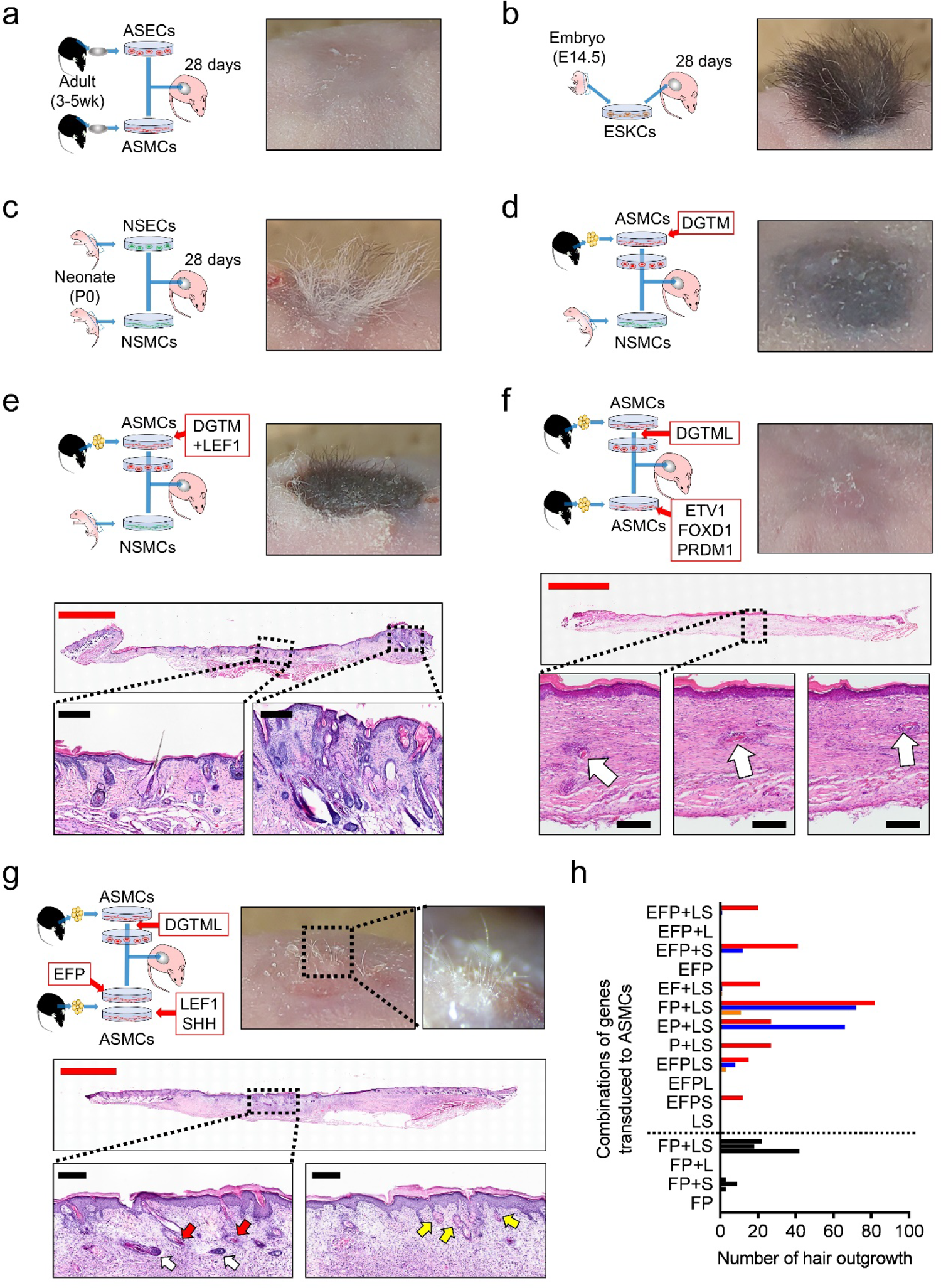
Identification of factors for induction of cells with the ability to form skin appendages. **a**, Experimental design schematic and representative image of reconstituted skin after transplantation of ASECs and ASMCs into a skin chamber on the back of immunodeficient mice. No skin appendage regeneration was confirmed. n = 3, all similar results. **b**, Transplantation of ESKCs robustly regenerated skin appendages. n = 3, all similar results. **c**, Transplantation of NSECs and NSMCs moderately regenerated skin appendages. n = 3, all similar results. **d**, Transplantation of DGTM-iSEPs and NSMCs did not regenerate skin appendages. n = 3, all similar results. **e**, Transplantation of DGTML-iSEPs and NSMCs regenerated skin appendages. Haematoxylin and eosin (H&E) staining of sections through reconstituted areas of skin, showing hair follicles and sebaceous glands, similar to those of surrounding skin, in the central portion. Red scale bar, 2 mm; black scale bars, 200 µm. n = 3, all similar results. **f**, Transplantation of DGTML-iSEPs and ASMCs transduced with *ETV1*, *FOXD1*, and *PRDM1*. A buried hair-follicle-like structure was histologically confirmed in the central portion. Arrows in serial magnified panels indicate mature hair shafts. Red scale bar, 2 mm; black scale bars, 200 µm. n = 1. **g**, Transplantation of DGTML-iSEPs with two AMSCs transduced with *ETV1*, *FOXD1* and *PRDM1,* and *LEF1* and *SHH*, respectively, results in hair outgrowth. Histologically, skin appendages were confirmed in the central portion. Red arrows indicate mature hair shaft, white arrows indicate dermal papilla, and yellow arrows in the serial magnified panel indicate sebaceous glands. Red scale bar, 2 mm; black scale bars, 200 µm. n = 3, all similar results. **h**, Amount of hair outgrowth after transplantation of DGTML-iSEPs and ASMCs transduced with combinations of genes (E, *ETV1*; F, *FOXD1*; P, *PRDM1*; L, *LEF1*; S, *SHH*). Different colour bars represent different series.

### Reprogramming factors *in vitro*

First, we aimed to determine the genes that reprogram ASMCs to epithelial cells capable of reconstituting skin appendages with the help of NMSCs. The DGTM factors were tested first and were found to be insufficient for ASMCs to regenerate skin appendages (Fig. 1d). To confer developmental epithelial characteristics, *LEF1*, *FOXD1*, and *SHH* were tested as additional factors^2, 12, 13^. Epithelial cells were successfully induced with DGTM factors plus *LEF1* or *FOXD1*, while the addition of *SHH* prevented the induction of epithelial cells. Using a skin reconstitution assay, DGTML-iSEPs (DGTM+*LEF1*-induced stratified epithelium progenitors) were found to reconstitute skin appendages together with NSMCs (Fig. 1e). To determine the set of genes required for reprogramming adult mesenchymal cells to mesenchymal cells with the capacity for skin appendage reconstitution, ASMCs were processed and co-transplanted with DGTML-iSEPs. Twenty fetal skin-associated genes dermal fibroblasts (hDFs) and measuring the levels of the dermal papilla markers *PROM1*, *CRABP1*, and *VCAN* (Extended Data Fig. 1a). Eligible genes were further tested by measuring the alkaline phosphatase (ALP) expression on transduction of each gene singly and in combination (Extended Data Fig. 1b, c). Trial transplantations were done in parallel with no positive findings. However, with the addition of *PRDM1*^15, 16^ to one of the candidate combinations, *ETV1* and *FOXD1*, hair-follicle-like structures were found beneath the skin in the centre of the reconstituted area (Fig. 1f). To further approximate the local environment of developing skin, AMSCs transduced with *LEF1* and *SHH* were additionally transplanted as a third cell type^17, 18^. Consequently, hair outgrowth was observed from a central location. A mature hair shaft, hair follicle, and structures similar to a sebaceous gland were confirmed histologically (Fig. 1g). To minimize the mesenchymal elements in terms of cell types and gene numbers, DGTML-iSEPs were transplanted with differential mesenchymal cells transduced with combinations of genes including *ETV1*, *FOXD1*, *PRDM1*, *LEF1*, and *SHH*, and the amount of hair outgrowth was evaluated. Despite substantial differences between the various combinations, the results indicated that the combination of *FOXD1*+*PRDM1*-transduced mesenchymal cells (FP-MCs) and *LEF1*+*SHH*-transduced mesenchymal cells (LS-MCs) conferred the

### Contribution of reprogrammed cells

To reveal the contribution of each cell type to skin appendages, DGTML-iSEPs, FP-MCs, and LS-MCs generated from GFP-mouse-derived ASMCs were co-transplanted with other unlabelled cells. In DGTML-iSEPs from GFP animals, the reconstituted area was identified as an uninterrupted GFP-positive surface with a clear border. The whole epithelial portion, including the skin appendages, consisted of GFP-positive DGTML-iSEPs (Fig. 2a). In FP-MCs or LS-MCs from GFP animals, the GFP-positive cells comprised the subcutaneous portion of the reconstituted skin area. ALP-positive cells localized in dermal papilla positions were partially GFP positive in both FP-MCs (Fig. 2b) and LS-MCs from labelled animals (Fig. 2c). To further elucidate the role of cells from the recipient animal, non-labelled DGTML-iSEPs, FP-MCs, and LS-MCs were transplanted to GFP nude mice. GFP-positive recipient animal-derived cells could be found in the subcutaneous portion of the reconstituted skin area, but could not be detected in skin appendages (Fig. 2d). Thus, it was demonstrated that the core elements of regenerated skin appendages consisted of DGTML-iSEPs, FP-MCs, and LS-MCs. FP, and LS factors could induce *de novo* generation of skin with appendages, we employed an assay of isolated skin ulcers. We surgically removed skin from the back of mice to generate an ulcer and isolated the resulting wound from the surrounding skin using a skin chamber sutured to the deep fascia (Fig. 3a). Isolated wounds did not close in the absence of treatment since the migration of epithelial cells into the wound and contraction of the surrounding skin was prevented^7^.

**Fig. 2:**
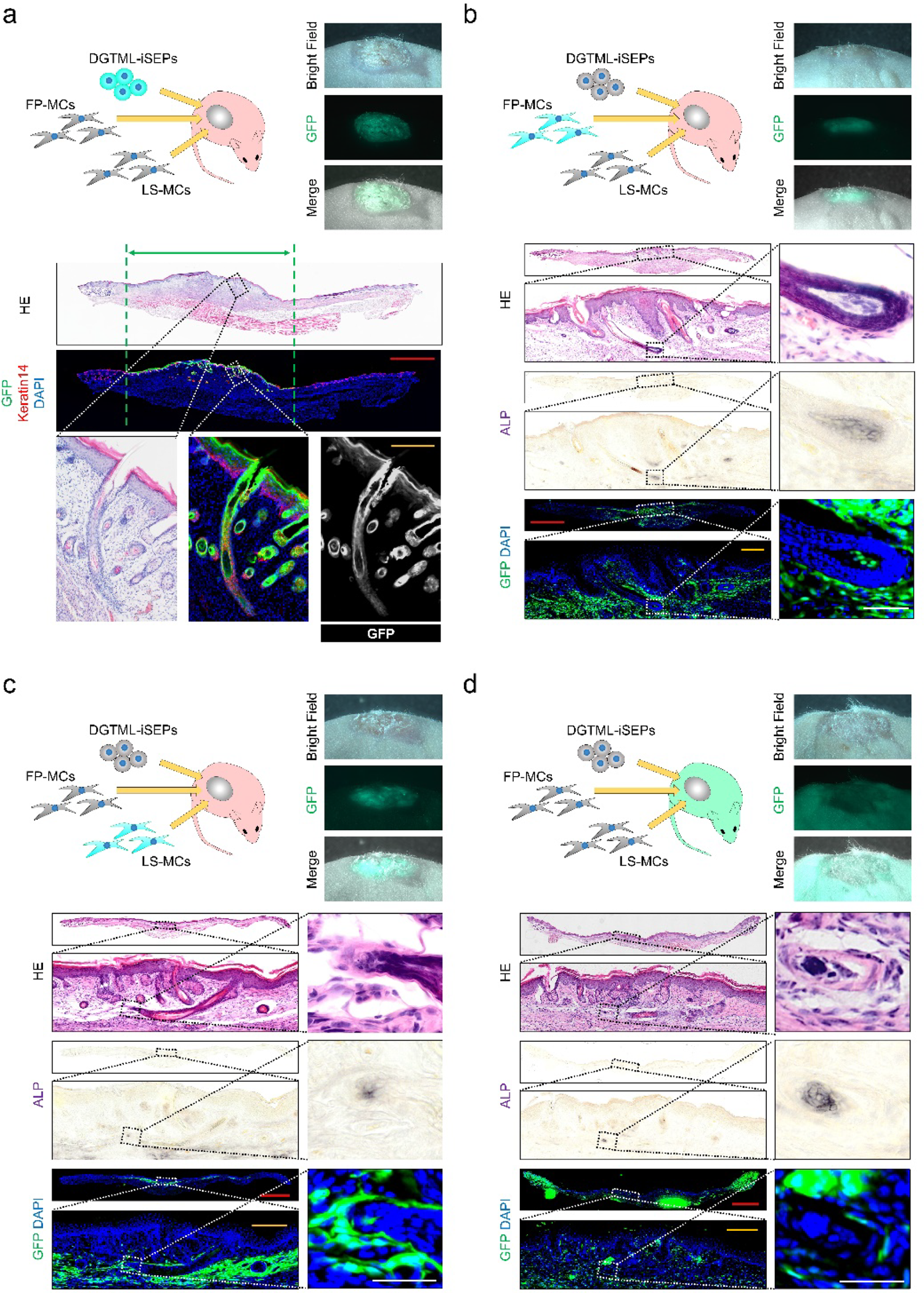
Induced cells contribute to skin appendages. **a**, Experimental design schematic and representative stereoscopic and histological findings of DGTML-iSEP tracing experiments. The green arrow indicates the area of GFP-positive epithelial surface. Images are representative of three independent experiments. **b**, Experimental design schematic and representative stereoscopic and histological findings of FP-MC tracing experiments. FP-MCs reside in the ALP-positive dermal-papilla-positioned cell cluster. Images are representative of nine independent experiments. **c**, Experimental design schematic and representative stereoscopic and histological findings of LS-MC tracing experiments. FP-MCs reside in the ALP-positive dermal-papilla-positioned cell cluster. Images are representative of nine independent experiments. **d**, Experimental design schematic and representative stereoscopic and histological findings of recipient animal-derived cell tracing experiments. Recipient animal-derived cells were not detected in skin appendages. Images are representative of six independent experiments. **a–d**, Haematoxylin and eosin (HE), alkali phosphatase (ALP), and immunofluorescent images were obtained from the same specimen slice. Red scale bars, 2 mm; orange scale bars, 200 µm; white scale bars, 50 µm.

**Fig. 3:**
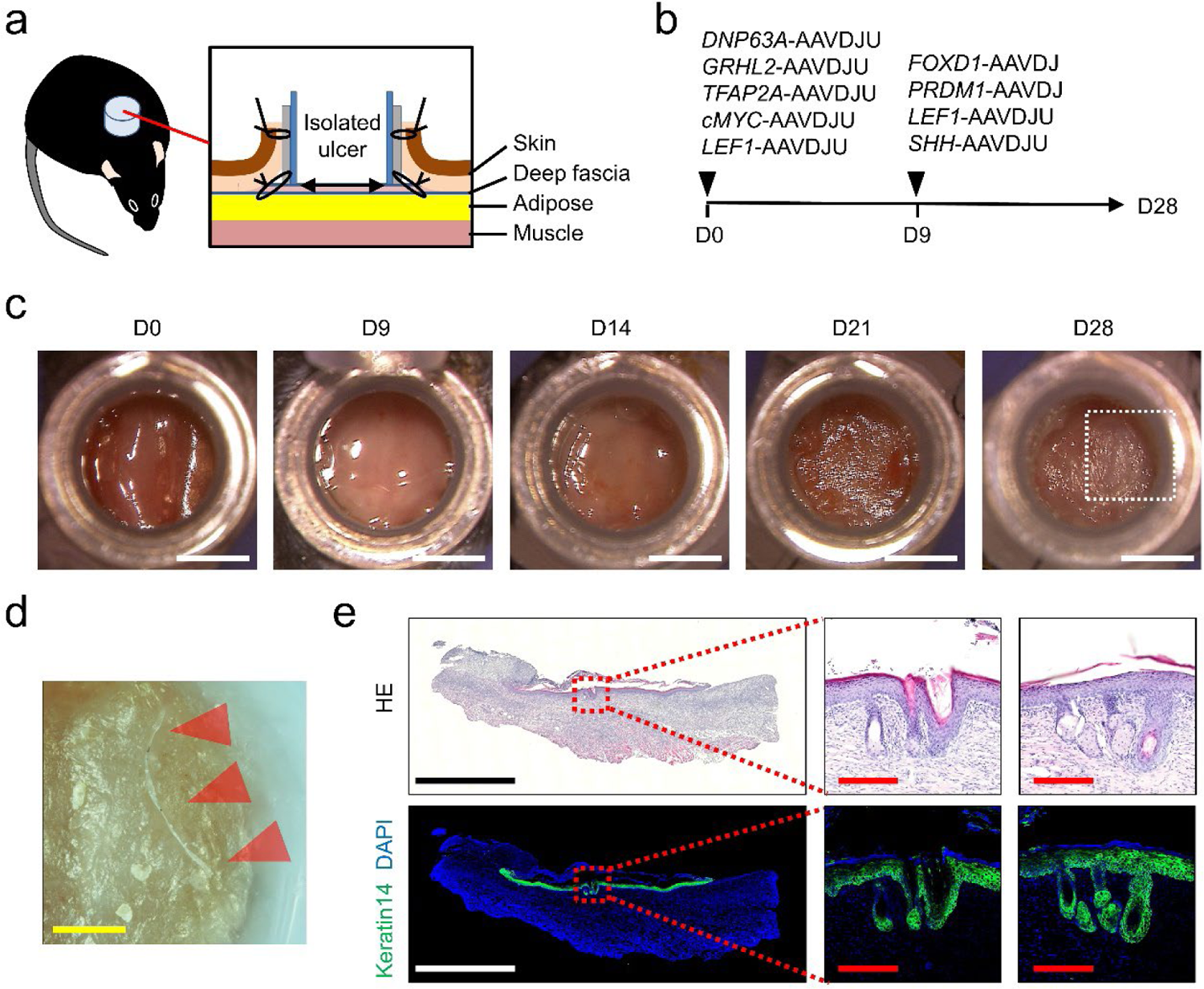
*In vivo* reprogramming of wound-resident cells to developing skin cells generates skin with appendages. **a**, Schematic of the skin chamber used for separating the ulcer from surrounding skin. **b**, Schedule of administration of reprogramming factors using two types of adeno-associated viruses (AAVs). **c**, Representative appearances of ulcers treated by AAVs. Epithelial-like structures were observed after day 21. Scale bars, 3 mm. (Similar findings in 1 of 4 animals in three independent experiments). **d**, Stereoscopic image of epithelial surface. Arrows indicate outgrowth of a hair-like structure. Scale bars, 1 mm. **e**, Haematoxylin and eosin (H&E) and immunohistochemical images of the same section through the root of the hair-like structure. A hair-follicle-like structure (middle panels) and a sebaceous-gland-like structure (right panels) were observed in isolated epithelia. Black-and-white scale bars, 2 mm; red scale bars, 200 µm.

### Development of a new AAV capsid

For *in vivo* gene transduction, we tested 18 wildtype and synthetic adeno-associated viral (AAV) capsids to analyse the cell type tropism between AAVs, and determined that AAVDJ (a serotype of the AAV capsid generated by DNA family shuffling^19^) was the most efficient for transducing cells in isolated ulcers. To develop an AAV capsid with increased efficiency and modified cell tropism, we applied *in vivo* directed evolution on an AAVDJ random peptide display library^20^ and generated a new AAVDJ-variant capsid optimized for cells in isolated murine skin ulcers (AAVDJU) (Extended Data Fig. 2a–e). The AAVDJU virus was more efficient than the original AAVDJ virus when applied in Data Fig. 3a–c).

### Reprogramming *in vivo*

We next applied DGTML-AAVDJUs (*DNP63A*-AAVDJU, *GRHL2*-AAVDJU, *TFAP2A*-AAVDJU, *cMYC*-AAVDJU, and *LEF1*-AAVDJU), followed by mixtures of FP-AAVDJs (*FOXD1*-AAVDJ and *PRDM1*-AAVDJ) and LS-AAVDJUs (*LEF1*-AAVDJU and *SHH*-AAVDJU) in our *in vivo* ulcer assay (Fig. 3b). We observed epithelia-like tissue inside the chamber around day 21 (Fig. 3c) and an outgrowth of a hair-like structure on day 28 (Fig. 3d) in one out of four animals (25%) in three independent series of experiments. Structures similar to hair follicles and sebaceous glands were confirmed in isolated epithelia on histological analysis (Fig. 3e). In summary, direct reprogramming of wound-resident cells to multiple developing skin cells enables the generation of structures similar to skin appendages *in situ*. The emergence of skin appendages from an isolated wound, even without epithelial components, supports the feasibility of *de novo* complex organ generation *in vivo*. These findings pave the way toward new therapeutics in which all regions of the wound re-observed during normal healing. An alternative supply of regenerative cells *in vitro* and *in vivo* might provide a new approach to frequent aging-associated skin appendage disorders, such as hair loss and dry skin. Our observations constitute an initial proof of principle for *in vivo* regeneration of three-dimensional complex tissue in mice. This knowledge might not only be useful for enhancing skin repair, but could also serve to guide *in vivo* regenerative strategies in other human pathological conditions in which tissue or organ homeostasis and repair are impaired.

## MATERIALS & METHODS

### Isolation and culture of mouse adult skin-derived epithelial cells (ASECs)

Back skin specimens were harvested from 3–5-week-old mice. The superficial portion was collected in strip form with scissors, and was incubated with 0.25% trypsin and 0.02% ethylenediaminetetraacetic acid (EDTA) in PBS for 16–24 hours at 4°C. The epidermis was separated from the dermis with forceps, and ASECs were isolated from the dermis. ASCEs were maintained on mitomycin C-treated 3T3-J2 feeder cells (a generous gift from the late Dr Howard Green) in F medium (3:1 [v/v] Ham’s F12 nutrient mixture:DMEM, high glucose (both from Life Technologies) supplemented with 5% FBS, 0.4 µg/ml hydrocortisone (Sigma), 5 µg/ml insulin (Sigma), 8.4 ng/ml cholera toxin (Wako), 10 ng/ml EGF (Wako), 24 µg/ml adenine (Sigma), 100 U/ml penicillin, 100 µg/ml streptomycin (Gibco), and 10 µM Rho-kinase inhibitor Y27632 (Selleck)^7^).

### Isolation and culture of mouse adult subcutaneous mesenchymal cells (ASMCs)

Subcutaneous groin-lumber fat pads were harvested from euthanized 3–5-week-old mice. Briefly, adipose tissue was enzymatically digested (as described^7^) and subsequently the stromal vascular fraction was isolated by centrifugation and inoculated on a gelatin-coated 6-well plate using one well for each mouse specimen, and maintained in complete DMEM growth medium, consisting of DMEM (containing 4.5 g/L glucose, 110 mg/L sodium pyruvate, and 4 mM L-glutamine) supplemented with 10% (v/v) heat-inactivated fetal bovine serum, 1:100 [v/v] MEM non-essential amino acid solution (Gibco), and 1:100 [v/v] GlutaMAX supplement (Gibco).

### Isolation and culture of mouse neonatal skin epithelial cells (NSECs) and neonatal skin mesenchymal cells (NSMCs)

Dorsal skin pieces were harvested from euthanized newborn mice (P0). Skin sheets were incubated with 0.25% trypsin and 0.02% EDTA for 16–24 hours at 4°C. In F medium, the epidermis was peeled off from the dermis using forceps. The resident epidermal cells were scraped from the dermis and epidermal cells and inoculated on mitomycin C-treated 3T3-J2 feeder cells in F medium, while dermal tissue was cut into pieces using a razor blade, digested with collagenase, and inoculated with complete DMEM growth medium in a gelatin-coated 6-well plate using one well for each mouse specimen.

### Isolation and culture of mouse embryonic skin cells (ESKCs)

After testing E12.5–E14.5 embryos, E14.5 was selected for use. Dorsal skin pieces were harvested from a euthanized mouse E14.5 embryo and incubated with 0.25% trypsin and 0.02% EDTA for 16–24 hours at 4°C. The sample was cut into pieces using a razor blade, digested with collagenase, and inoculated with F medium in a gelatin-coated 6-well plate, using one well for each mouse specimen.

### Mouse iSEP generation

Primary-culture ASMCs of more than two passages were seeded at 20,000 cells per well in 24-well culture plates. The next day, AAVs (1.0×10^9^–10^10^) were mixed with complete DMEM medium. The medium was changed on days 1, 2, and 4. The medium was changed to F medium from day 4 or 5. The medium was changed daily. After the emergence of epithelial colonies, cells were passaged and maintained with the same protocols as ASECs.

### Culture of human dermal fibroblasts (hDFs)

Normal human dermal fibroblasts (Cat. # C-12300) were purchased from PromoCell (Heidelberg, Germany) and maintained in complete DMEM growth medium.

### Retroviral plasmid construction

candidate factors were prepared by subcloning the ORF template clones by PCR amplification with Prime STAR GXL DNA polymerase (Takara) and ligation by In-Fusion cloning enzyme (Clontech).

### Retrovirus production

For retrovirus production, pMXs vectors were co-transfected with packaging plasmids (pCMV-gagpol-PA and pCMV-VSVg) into 293FT cells (Thermo Fisher Scientific) using Lipofectamine 2000 (Thermo Fisher Scientific). Retroviral supernatants were collected 48 hours after transfection and debris was excluded by centrifugation twice for 20 minutes at 2000 × *g*.

### AAV plasmid construction

AAV plasmids were prepared by subcloning the ORF to pAAV-CAG-DNP63A^7^ or using the In-Fusion HD Cloning kit (Clontech).

### AAV production

AAVs were prepared using 293AAV cells (Cell Biolabs, Inc.) by calcium phosphate titre was determined by qPCR using the primers ITR-F, 5′-GGAACCCCTAGTGATGGAGTT-3′ and ITR-R, 5′-CGGCCTCAGTGAGCGA-3′.

### AAVDJ peptide display library

The backbone plasmid for cloning the random oligonucleotides was generated from pAAV-CAG-GFP (Plasmid #37825, Addgene) and pAAVDJ using the In-Fusion HD Cloning kit (Clontech) and QuikChange II Site-Directed Mutagenesis Kit (Agilent), with reference to previous reports^19, 20^. The first-round peptide display library plasmid was prepared using an *Sfi*I-digested backbone and a *Bgl*I-digested random-trimer oligonucleotide (Ella Biotech GmbH) using T4 DNA ligase (New England Biolabs). The AAVDJ peptide display library virus was produced by transfection of reduced library plasmid (1% (w/w)) with helper plasmids.

### qPCR analyses for dermal papilla markers

Human DFs were seeded at 30%–40% confluency in 12-well culture plates. The next day, retroviral-containing supernatant (up to 40% of total medium) was mixed with complete changed on days 1 and 2. On day 4, total mRNA was purified (ZYMO Research, CA, R1058, Quick-RNA MINIprep plus), reverse transcribed (Thermo Fisher Scientific, M1662, Maxima™ H Minus cDNA Synthesis Master Mix), and analysed with qPCR (TOYOBO Bio, QPS-101, THUNDERBIRD® qPCR Mix) for *PROM1*, *CRABP1*, and *VCAN* using the following primers: *PROM1*-F, 5′-GGACCCATTGGCATTCTC-3′ and *PROM1*-R, 5′-CAGGACACAGCATAGAATAATC-3′; *CRABP1*-F, 5′-GCAGCAGCGAGAATTTCGAC-3′ and *CRABP1*-R, 5′-CGTGGTGGATGTCTTGATGTAGA-3′; *VCAN*-F, 5′-GTAACCCATGCGCTACATAAAGT-3′ and *VCAN*-R, 5′-GGCAAAGTAGGCATCGTTGAAA-3′; and *GAPDH*-F, 5′-GGAGCGAGATCCCTCCAAAAT-3′ and *GAPDH*-R, 5′-GGCTGTTGTCATACTTCTCATGG-3′.

### Alkaline phosphatase assay

Four days after retroviral transfection to hDFs (as done for qPCR analyses), cells were fixed and stained using Stemgent® Alkaline Phosphatase Staining Kit II (Reprocell, Japan). The number of alkaline phosphatase (ALP)-positive cells in each well (24-well stereoscopy and analysed for positive cell counts using ImageJ.

### *In vivo* biopanning in ulcers

The AAVDJ peptide display library virus was inoculated into an isolated skin ulcer in a silicone chamber. After 2–4 days, cells in the ulcer were isolated from tissues above the thoracic wall by collagenase digestion and inoculated on a gelatin-coated 6-well plate, as for ASMCs. After 2 days, genomic DNA was purified from cells using a DNeasy Blood & Tissue Kit (QIAGEN). Randomized capsid sequences were PCR amplified. *Hin*dIII/*Not*I-digested PCR reactant and backbone were ligated by T4 DNA ligase (New England Biolabs) and transformed to CloneCatcher™ (Genlantis) using ELEPO21 (Nepagene). The next-round AAVDJ library was produced by transfection of a reduced library plasmid (1% (w/w)) with helper plasmids.

### AAVDJ capsid engineering

Six *in vivo* biopanning cycles were applied for four series of animals. Twelve clones for each series were sequenced by Sanger sequencing. Amino acid sequences confirmed in two series of experiments were selected as expected variants, PCR amplified, and evaluated for 10^10^ (gene copies (GC) virus/well) GFPNLS (green fluorescent protein (GFP) with a nuclear localization signal (NLS))-expressing engineered AAVs in 24-well plates.

### Comparative analyses of AAVDJ and AAVDJU

GFPNLS- and mCherry-expressing AAVDJ (original AAVDJ) virus and AAVDJU (engineered AAVDJ) virus were prepared. We inoculated 50 µl of virus solution, including 10^11^ GC of GFPNLS-expressing virus and 10^11^ GC of mCherryNLS-expressing virus (i.e. AAVDJ-GFPNLS + AAVDJ-mCherryNLS, AAVDJU-GFPNLS + AAVDJU-mCherryNLS, AAVDJ-GFPNLS + AAVDJU-mCherryNLS, and AAVDJU-GFPNL + AAVDJ-mCherryNLS), to ulcers in silicone chambers attached on the interscapular area of mice (n = 5 for each group). Four days later, the top of the chamber was cut off and the ulcer surface was imaged using stereoscopy. The chamber and surrounding tissues were collected, fixed, and embedded in OCT compound. For each sample, sections (more than 400 µm apart) through the isolated skin ulcer were prepared and analysed to calculate the GFPNLS- and mCherryNLS-expressing cell frequency.

### Histological analyses of gene transduction efficiency

With a guidance from the HE-stained serial section, the area of the DAPI-stained section was classified as the superficial layer (above the fascia of subcutaneous adipose tissue), the adipose layer (subcutaneous adipose tissue), and the muscle layer (striated muscle, such as trapezius muscle). To segment the nuclei in each layer, we generated UNet++ models trained with the RMSprop optimizer and binary cross-entropy dice coefficient loss (BCE-dice-loss). The images for generating the training data were cropped to 512 × 512 pixels and subjected to model training with 400 epochs, a batch size of 8, and a learning rate of 10^−4^. After the training, the model was applied to all the images to predict the nuclear area, and the probability of each pixel was calculated in the range of [0, 1]. Pixels with probabilities in the range of [0.5, 1] were annotated as nuclei, and then the nuclear regions were segmented using the watershed algorithm. Nuclei with more than 5 GFP-positive pixels were defined as nuclei from AAV-infected cells. The numbers of AAV-infected nuclei were counted for each region and layer. For each sample, 10–16 sections (mean, 14.4) were analysed.

### Skin reconstitution assay

BALB/cAJcl-nu/nu female mice were used as recipient animals. Under inhalation autoclaved 1.0-cm diameter silicone chamber, generated using a 3D printed template^11^, was inserted into the skin hole. Four 5-0 nylon sutures were made to attach the rim of the silicone chamber to the skin. Epithelial cells and mesenchymal cells (1–6 wells in a 6-well plate), for investigation of the skin appendage regeneration ability, were prepared in 150 µl 1:1 (v/v) mixtures of keratinocyte F medium and complete DMEM growth medium and transferred into the chamber via a small incision made at the top of the chamber. On day 7, the upper half of the chamber was cut off. On day 14, the chamber was removed. The regeneration of skin appendages was evaluated on day 28.

### Assay for identification of reprogramming factors

Epithelial cells to be assessed for skin appendage regeneration ability were transplanted after purification by cultivation with NMSCs passaged more than twice (early passage NMSCs include NSECs, resulting in false-positive skin appendage regeneration). Mesenchymal cells to be assessed were transplanted with DGTML-iSEPs 4–8 days after the transduction of retroviral candidate genes.

### *In vivo* skin regeneration assay

ulcers, we aimed to induce epithelial tissues in skin ulcers that were isolated from the surrounding skin by a skin chamber in female C57BL/6JJcl mice. To avoid subcutaneous glands, such as the thyroid and mammary glands, the interscapular area was chosen as the optimal site for chamber attachment. Chambers were made by cutting 0.6-ml low-attachment microtubes (BM4006 from BM Bio). The lateral surface of the tubes was covered by a silicon tube with an external diameter of 6 mm and an internal diameter of 5 mm for suture fixation. Under inhalation anaesthesia, the chamber attachment site was shaved and sterilized. From a 1.5 cm vertical incision on the spine, a flap was elevated beneath the panniculus carnosus, and the chamber was sutured down to the deep fascia with six horizontal mattress sutures and sutured to the surrounding skin flaps to ensure fixation/sealing of the chamber using 4-0 Ethilon (Ethicon Inc.). Then, AAVs were administered and the lid was closed. The edge of the skin flap was glued to the chamber when necessary.

### Animals

C57BL/6JJcl and BALB/cAJcl-nu/nu mice were purchased from Nippon Bio-Supp. Center. C57BL/6-Tg(CAG-EGFP) and C57BL/6-BALB/c-nu/nu-EGFP mice were indicated. All animal procedures were approved by the IACUC of the Graduate School of Medicine and Faculty of Medicine, The University of Tokyo.

### Histological analyses

For frozen sectioning, samples were fixed using 4% paraformaldehyde in PBS, then incubated in 30% sucrose in PBS for 1–2 days. Tissues were embedded in OCT compound and were frozen using dry ice. Tissue sections were routinely stained with haematoxylin and eosin. Immunohistochemical analyses were performed using antibodies against Cytokeratin 14 (ab181595, 1:500; Abcam). Secondary antibodies were obtained from Life Technologies. After staining, sections were mounted with DAPI Fluoromount-G (Southern Biotech). Sections processed for confirmation of the cell contribution to regenerated skin appendages were first stained with BCIP®/NBT solution (Sigma B6404) and DAPI (Dojindo D523, Kumamoto, Japan, 1:1,000), analysed, then stained with haematoxylin and eosin and re-analysed. Sections analysed for *in vivo* generation of skin appendages were immunohistochemically analysed first, then stained with haematoxylin and eosin and re-analysed.

### Imaging analyses

Imaging analyses were performed using a Confocal Zeiss LSM900, a Stereoscope Zeiss AXIO Zoom.V16, an Olympus VS-200 Slide Scanner, an Olympus IX73 Inverted LED Fluorescence Microscope, and a high-resolution operative microscope Mitaka Kohki MM100-YOH.

### Statistical analysis

No statistical methods were used to predetermine sample size. The experiments were not randomized, and the investigators were not blinded to allocation during experiments and outcome assessment.

### Reporting summary

Further information on research design is available in the Nature Research Reporting Summary linked to this paper.

## Acknowledgements

This work was supported by JSPS KAKENHI Grant numbers JP20H03847 (to M. K.); JP20K20609 (to M. K.); 19H03813 (to M. O.); 22H03247 (to M. O.); AMED under Grant Catherine Perfect, MA (Cantab), from Edanz (https://jp.edanz.com/ac), for editing a draft of this manuscript.

## Author Contributions

M. Ku. conceived the project. M. Ku. planned the experiments. Y. M., S. Q., D. Z., M. Ka., T. N., K. L., K. K., and M. Ku. performed the experiments. H. O., S. S., K. K., Y. H., H. H., and M. Ku. analysed the results, and M. O., J. C. I. B., and M. Ku. wrote the manuscript with editing by all the other authors.

---

**Extended Data Fig. 1.**
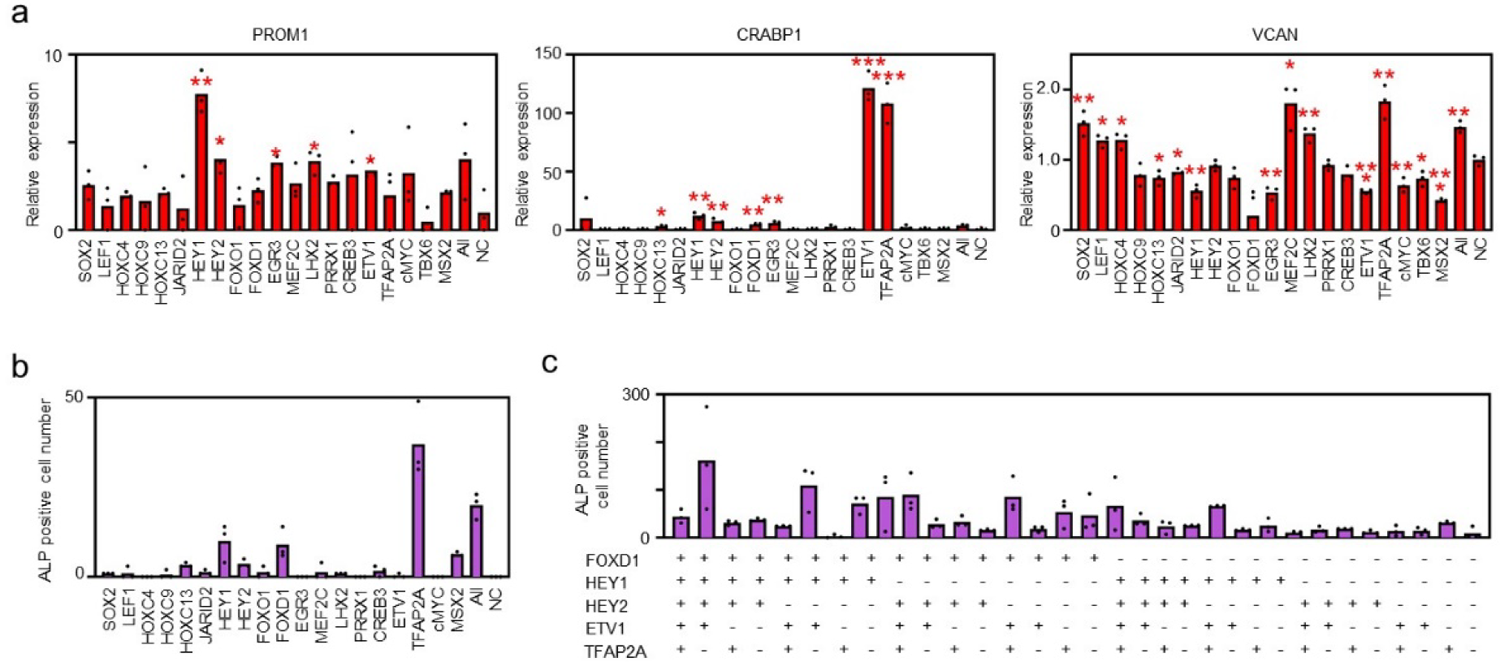
**a**, Changes in expression levels of the dermal papilla markers *PROM1*, *CRABP1*, and *VCAN* after candidate gene transduction as assessed by quantitative PCR. **b**, Number of alkaline phosphatase (ALP)-positive cells in each well (24-well plate) four days after candidate gene transduction. Overlaid dot plots indicate the distribution of the data (n = 3, technical replicates). **c**, Number of ALP-positive cells in the central field of 24 wells after candidate gene transduction. Overlaid dot plots indicate the distribution of the data (n = 3, technical replicates).

**Extended Data Fig. 2:**
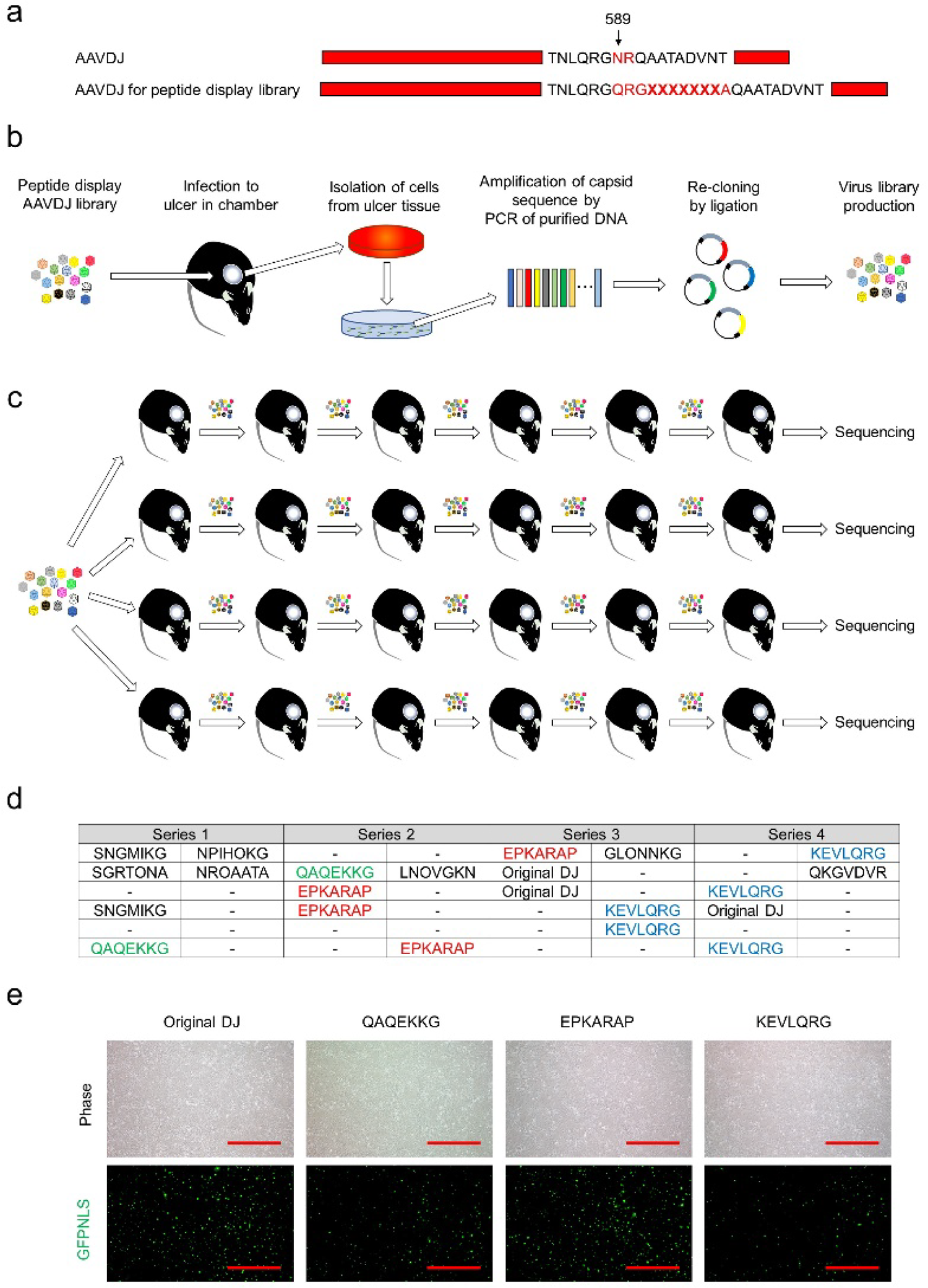
Development of an AAVDJ-variant capsid optimized for cells in murine isolated skin ulcers. **a**, Schematics of the AAVDJ peptide display library. The library was generated by mutagenesis of AAVDJ at the 589^th^ amino acid position. X, randomized amino acid. **b**, Schematics of the *in vivo* biopanning cycle. A library virus was inoculated in an isolated skin ulcer in a chamber. After 2–4 days, cells were isolated from the ulcer tissue. After two days, genomic DNA was purified from the cells. Randomized capsid sequences were PCR amplified and subcloned to a backbone vector. The next-cycle virus library was generated. **c**, Schematics of capsid engineering. Six *in vivo* biopanning cycles were applied for four series of animals. **d**, Randomized amino acid sequences detected by sequencing of 12 clones for each series. (-) indicates the sample was unreadable. Amino acid sequences highlighted in red, blue, and green were sequenced in two series of experiments and rendered for *in vitro* gene transduction analysis. **e**, Findings of ASMCs four days after infection with GFPNLS (green fluorescent protein (GFP) with a nuclear localization signal (NLS))-expressing original DJ virus and the DJ-variant virus harnessing selected amino acid sequences. The DJ variant with EPKARAP was more efficient than other two variants and thus was employed as a new AAVDJ-variant capsid optimized for cells in murine isolated skin ulcers (AAVDJU). Scale bar = 500 µm.

**Extended Data Fig 3:**
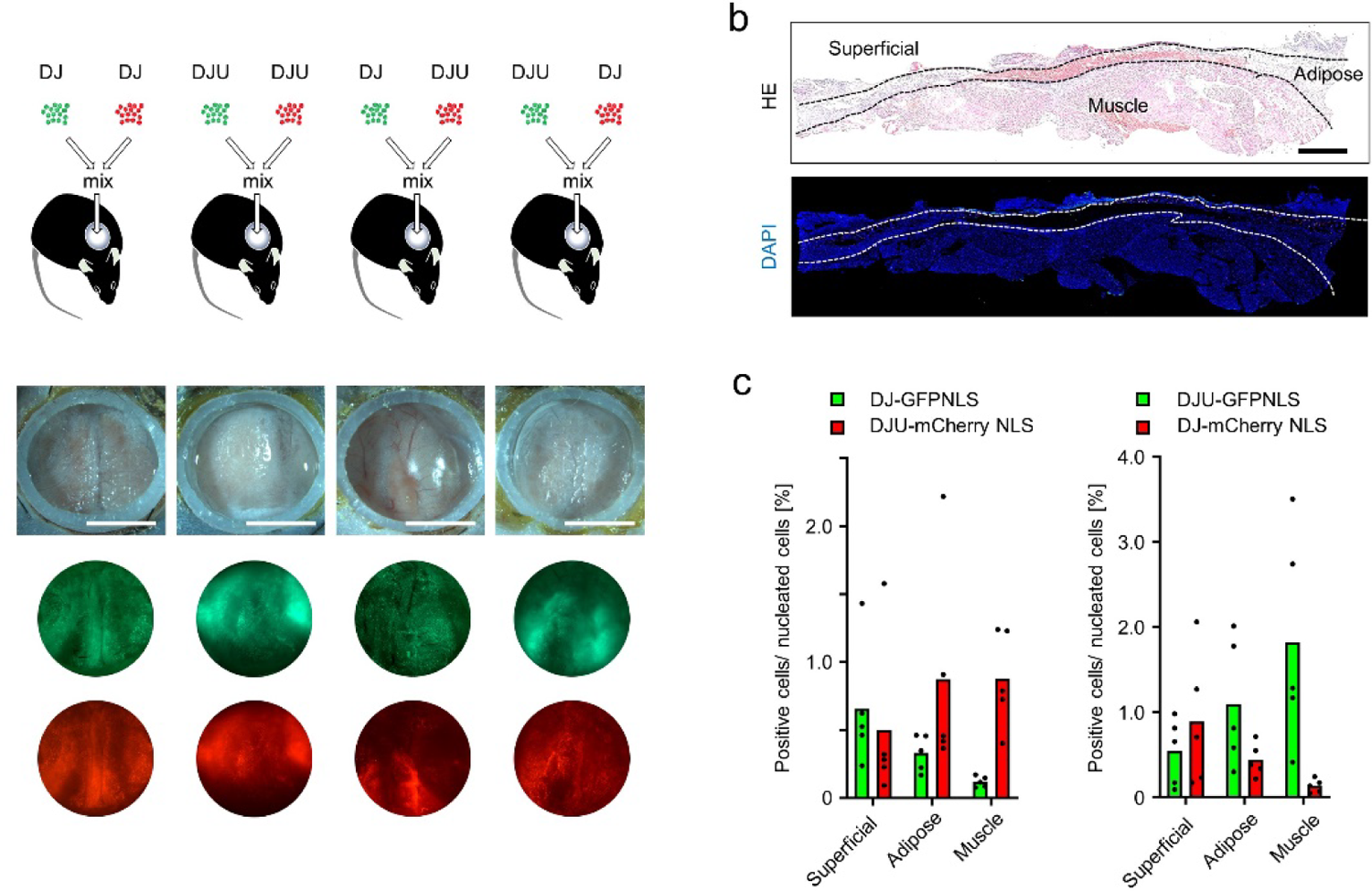
AAVDJU is superior to AAVDJ for cells in the deep layer of a murine isolated skin ulcer. **a**, (top) Schematics of comparative analyses of the gene transduction efficiency of AAVDJ and AAVDJU. AAVDJ-GFPNLS, AAVDJ-mCherryNLS, AAVDJ-GFPNLS, and AAVDJ-mCherryNLS were prepared, mixed, and inoculated in isolated ulcers (n = 5 for each condition). (2^nd^ row) Stereoscope images of representative animals four days after inoculation with AAVs. (3^rd^ and 4^th^ row) Stereoscope fluorescence images of the ulcer surface. The fluorescence expression patterns from the same capsid, such as AAVDJ-GFPNLS and AAVDJ-mCherryNLS, were similar, while those from different capsids were different. Samples were collected and rendered for histological analyses. Scale bar = 5 mm. **b**, Haematoxylin and eosin (H&E) staining and fluorescent images of a section through an isolated skin ulcer. The gene transduction efficiencies were analysed in fluorescent images for three tissue layers (superficial, adipose, and muscle) demarcated by neighbouring H&E images. Scale bar = 10 mm. **c**, Frequency of nuclear-fluorescent-signal-positive cells in each layer of tissue in animals to which AAVDJ-GFPNLS plus AAVDJU-mCherryNLS (n = 5) and AAVDJU-GFPNLS plus AAVDJ-mCherryNLS (n = 5) were delivered, respectively, histologically quantified by deep-learning assisted imaging analyses. In both groups of animals, AAVDJU was more efficient than AAVDJ in the adipose and muscle layers.

